# IgA Potentiates NETosis in Response to Viral Infection

**DOI:** 10.1101/2021.01.04.424830

**Authors:** Hannah D. Stacey, Diana Golubeva, Alyssa Posca, Jann C. Ang, Kyle E. Novakowski, Muhammad Atif Zahoor, Charu Kaushic, Ewa Cairns, Dawn M. E. Bowdish, Caitlin E. Mullarkey, Matthew S. Miller

**Author notes:** Corresponding author: McMaster University, 1280 Main St. W, Hamilton, ON, Canada, L8S 4K1. Toronto Center for Liver Disease, Toronto General Hospital Research, Institute, MaRS-Princess Margaret Cancer Research Tower 10-401, University Health Network, 101-College St. Toronto, ON, Canada.

## Abstract

IgA is the second most abundant antibody present in circulation and is enriched at mucosal surfaces. As such, IgA plays a key role in protection against a variety of mucosal pathogens, including viruses. In addition to neutralizing viruses directly, IgA can also stimulate Fc-dependent effector functions via engagement of Fc alpha receptors (FcαRI) expressed on the surface of certain immune effector cells. Neutrophils are the most abundant leukocyte, express FcαRI, and are often the first to respond to sites of injury and infection. Here, we describe a novel function for IgA:virus immune complexes (ICs) during viral infections. We show that IgA:virus ICs potentiate NETosis – the programmed cell death pathway through which neutrophils release neutrophil extracellular traps (NETs). Mechanistically, IgA:virus ICs potentiated a suicidal NETosis pathway via engagement of FcαRI on neutrophils through a toll-like receptor (TLR)-independent, NADPH oxidase complex-dependent pathway. NETs also were capable of trapping and inactivating viruses, consistent with an antiviral function.

## INTRODUCTION

IgA antibodies have pleiotropic roles in regulating the response to microbes. In the context of infection, IgA antibodies enriched at mucosal surfaces as secretory IgA (sIgA) are capable of neutralizing viruses in an “anti-inflammatory” manner since these antibodies block infection but do not activate immune cells via Fc receptor engagement. However, monomeric IgA (mIgA) antibodies, which are abundant in serum, are capable of engaging Fc receptors on the surface of immune cells to elicit effector functions ^1^.

Neutrophils are not only the most abundant leukocytes, but are often the first to respond to sites of injury and infection ^2^. Human neutrophils express the Fc alpha receptor (FcαRI/CD89) and are capable of exerting a variety of effector functions including phagocytosis, respiratory burst, antibody dependent cellular phagocytosis (ADCP), and NETosis ^3,4^. Data regarding the protective versus pathogenic role of neutrophils during viral infection is nuanced and suggests context is critical in determining outcome. For example, while neutrophils are required for protection during the early stages of influenza A virus (IAV) infection, neutrophils also release reactive oxygen species (ROS), proteolytic enzymes, and a variety of inflammatory mediators that can damage lung tissues. As a result, excessive neutrophil infiltration has been associated with severe lung injury ^5^.

The generation of NETs was first described by the Zychlinsky laboratory in 2004 as an antibacterial effector mechanism ^6^. NETs produced via a specialized form of programmed cell dealth called “NETosis” and are composed primarily of decondensed chromatin studded with antimicrobial proteins. Extensive work by many laboratories has since demonstrated that NETs can have not only protective, but also pathogenic consequences in infections and many other diseases ^4^. The understanding of how NETs influence viral infections continues to evolve. In the context of Chikungunya virus and poxvirus, NETs were capable of trapping virus and controlling infection in a mouse models of disease ^7,8^. Likewise, NETs have been shown to trap and inactivate HIV ^9^. However, NETs have also been described to exacerbate disease in the context of Dengue virus, rhinovirus, respiratory syncytial virus, influenza virus, and most recently, SARS-CoV-2 infection ^10–16^. Thus, the overall impact of NETs during a viral infection must be interpreted carefully in conjunction with other infection parameters.

Recently, Fc-dependent effector functions have been shown to play a central role in the protection conferred by broadly-neutralizing antibodies (bnAbs) that bind to the hemagglutinin (HA) stalk domain of IAV ^17–19^. However, these studies have only been performed in the context of monoclonal IgG antibodies. Elicitation of bnAbs is now the goal of several “universal” influenza virus vaccine candidates, including “chimeric” HA vaccines that were recently tested in a Phase I clinical trial ^20^. Antibody-dependent cellular cytotoxicity (ADCC) may also augment protection mediated by human immunodeficiency virus (HIV)-neutralizing antibodies ^21^. However, despite the fact that both IAV and HIV are mucosal pathogens – almost nothing is known about the contribution of IgA-mediated Fc-dependent effector functions during infection. This is due, in large part, to the fact that mice do not express an FcαR homolog which presents significant challenges for assessing the contributions of IgA to outcomes *in vivo* ^1^.

Here, we show that IgA:virus ICs potentiated NETosis through FcαRI signaling on neutrophils. This potentiation was not virus-specific, and could be observed for IAV, HIV, SARS-CoV-2 and extended to IgA ICs generated with antibodies/autoantigens from RA patients. In contrast to NETosis stimulated by virus directly, IgA:virus ICs stimulated suicidal NETosis that was independent of TLR signaling. Finally, viruses were trapped and inactivated in NETs, suggesting a protective role *in vivo* when properly regulated.

## RESULTS

### IgA:IAV immune complexes stimulate NETosis

Historically, antibodies have been thought to mediate protection against influenza viruses primarily by binding to the HA head domain and blocking interaction between the receptor binding site on HA and sialic acids on the surface of host cells. However, more recently it has become clear that bnAbs that bind to the HA stalk domain mediate protection *in vivo* primarily by elicitation of Fc-dependent effector functions ^17–19^. Antigen-specific IgA antibodies have been shown to neutralize IAV, but relatively little is known about IgA-mediated Fc-dependent effector functions during IAV infection ^22^. Neutrophils are the most abundant leukocyte and are among the first to respond during IAV infection ^23^. Neutrophils also express FcαRI, and we have previously shown that IgA:IAV ICs stimulate ROS production in neutrophils; however, unlike IgG:influenza virus ICs, this could not be fully inhibited by cytochalasin D – indicating that IgA-mediated ROS production was not due to antibody-dependent cellular phagocytosis (ADCP) ^24^. To determine whether IgA was capable of potentiating NETosis upon binding IAV, neutrophils were exposed to antibody:IAV ICs composed of polyclonal (monomeric) IgA or IgG from the peripheral blood of donors previously vaccinated with seasonal influenza vaccines containing the A/California/04/2009 (Cal/09) H1N1 component. Phorbol 12-myrisate13-acetate (PMA), a potent inducer of NETosis, was used as a positive control ^25^. IgA:IAV ICs stimulated significantly higher levels of NETosis than antibodies or virus alone, whereas IgG:IAV ICs did not induce NETosis above background levels (Fig. 1A, B).

**Figure 1.**
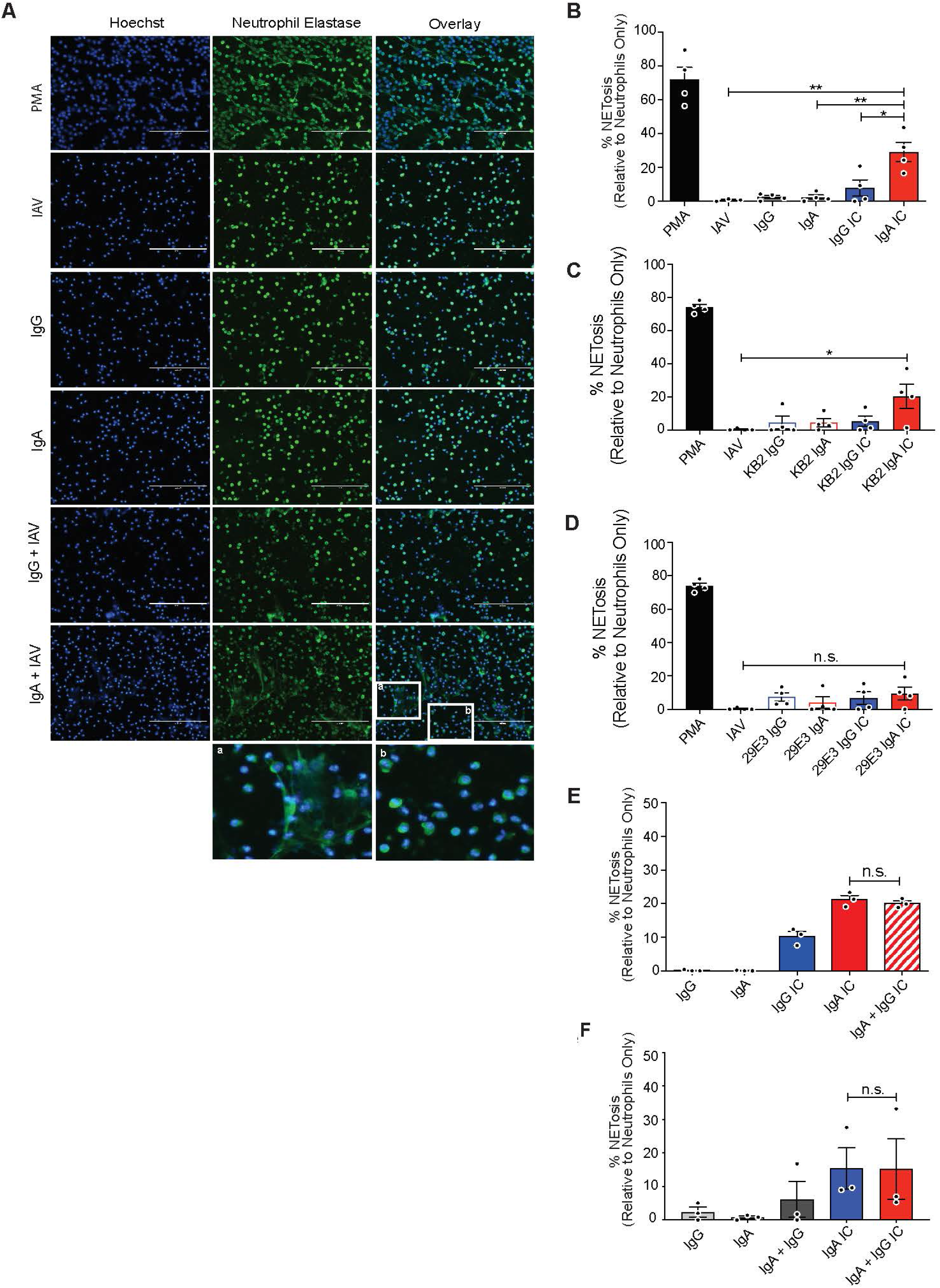
IgA:IAV ICs potentiate NETosis. (A-F) Primary human neutrophils were isolated from the peripheral blood of healthy donors (n=3 or 4), and stimulated with PMA, monoclonal or polyclonal IgG or IgA antibodies, or ICs for 3 hours as shown. NETosis was assessed by immunofluorescence microscopy after co-staining for DNA (DAPI) and neutrophil elastase. (A) Representative images are shown (20x). Bars depict 200 µm. Insert (a) shows area with NETs, insert (b) shows area with intact neutrophils. (B) The percentage of cells that had undergone NETosis (defined by typically NET morphology and co-staining of DAPI + neutrophil elastase) were quantified in a blinded manner from 5 fields in 4 independent experiments. (C, D) The assay was repeated using monoclonal antibodies, (C) KB2 and (D) 29E3, which bind the HA stalk and head domain of Cal/09, respectively. (E, F) To determine the phenotype of mixed IgG/IgA ICs, polyclonal IgG and IgA were mixed with Cal/09 at a (E) 1:1 or (F) at the ratio naturally found in serum. For all experiments, percent NETosis was normalized to unstimulated neutrophils. Three or four independent neutrophil donors were used for each experiment. Means and standard error (SEM) of independent experiments are shown. Statistical significance was determined using one-way ANOVA with Tukey post-hoc test. *,P < 0.05; **,P < 0.01.

In the context of IgG, bnAbs that bind to the stalk domain have been shown to potently elicit Fc-dependent effector functions, whereas antibodies that bind to the HA head domain and exhibit hemagglutination inhibiting (HAI) activity do not. This is because HA stalk-binding bnAbs allow for a two points of contact between target and effector cells ^26,27^. To determine whether broadly-neutralizing IgA:IAV ICs are primarily responsible for the induction of NETosis observed in the context of IAV-specific polyclonal IgA, we used a panel of previously-described monoclonal antibodies that bind to neutralizing epitopes on either the HA head or stalk domains ^28–30^. The antibody KB2 binds to the HA stalk domain of H1 viruses, while 29E3 is specific to the HA head domain of Cal/09 ^31^. When human neutrophils were incubated with ICs containing a IAV:IgA stalk-binding antibody (KB2), significant induction of NETosis was observed following 3-hour stimulation (Fig. 1C). In contrast, NETosis was not induced by IgG:IAV ICs, or by antibodies or virus alone (Fig. 1C). All ICs generated with a HA head-binding antibody (29E3) failed to induce NETosis (Fig. 1D).

In blood, IgA co-circulates with other antibodies, including IgG, which signals through distinct FcRs (FcγRs) and can also induce NETosis ^32^. Mixed ICs composed of IgG/IgA:HIV have also been shown to act cooperatively to stimulate ADCC by monocytes ^33^. We therefore tested whether mixed ICs composed of IAV bound by IgA and IgG together would influence the magnitude of NETosis induction relative to IgA alone. When ICs were generated with a 1:1 ratio of IgG:IgA, the magnitude of NETosis induction was similar to IgA alone (Fig. 1E). In serum, IgG is significantly more abundant than IgA (approx. 4:1 to 10:1). Thus, to recapitulate the physiological stoichiometry of IgG:IgA, we purified each immunoglobulin from serum of matched donors, and then recombined them at their natural physiological ratio. Here again, the magnitude of NETosis observed in mixed IgG:IgA immune complexes was similar to IgA alone, indicating that IgG does not potentiate IgA-mediated NETosis, nor does it interfere with the ability of IgA to stimulate NETosis (Fig. 1F). Taken together, these results demonstrate that IgA:virus ICs stimulate neutrophils to undergo NETosis.

### IgA-mediated NETosis is not an IAV-specific phenomenon

NETs have been observed in the context of many other infections, including those caused by SARS-CoV-2 and HIV. In these studies, viruses were presumed to stimulate NETosis directly ^14,16,34,35^. We thus performed an experiment to test the amount of virus needed to stimulate NETosis independent of FcαR signaling. Neutrophils were stimulated with increasing concentrations of purified lentiviruses pseudotyped with the SARS-CoV-2 spike protein. A significant elevation in NETosis was observed when neutrophils were exposed to 0.05 and 0.2 mg/mL of purified virus (Fig. 2A). We then purified IgA from convalescence serum of a SARS-CoV-2 infected individual, and a SARS-CoV-2 naïve individual, incubated them with sub-stimulatory concentrations (0.0125 mg/mL) of spike pseudotyped lentiviruses to allow for IC formation, and then incubated these mixtures with primary human neutrophils from healthy donors. As we observed in the context of IAV, IgA:virus IC generated with IgA purified from SARS-CoV-2 convalescence serum was capable of stimulating NETosis, whereas pseudovirus:IgA mixtures from naïve serum was not (Fig 2B). These results confirm that IgA:virus ICs more potently stimulate NETosis when compared to virus alone, and that ICs are required for this potentiation, since IgA from seronegative individuals did not significantly induce NETosis when mixed with pseudotyped lentivirus.

**Figure 2.**
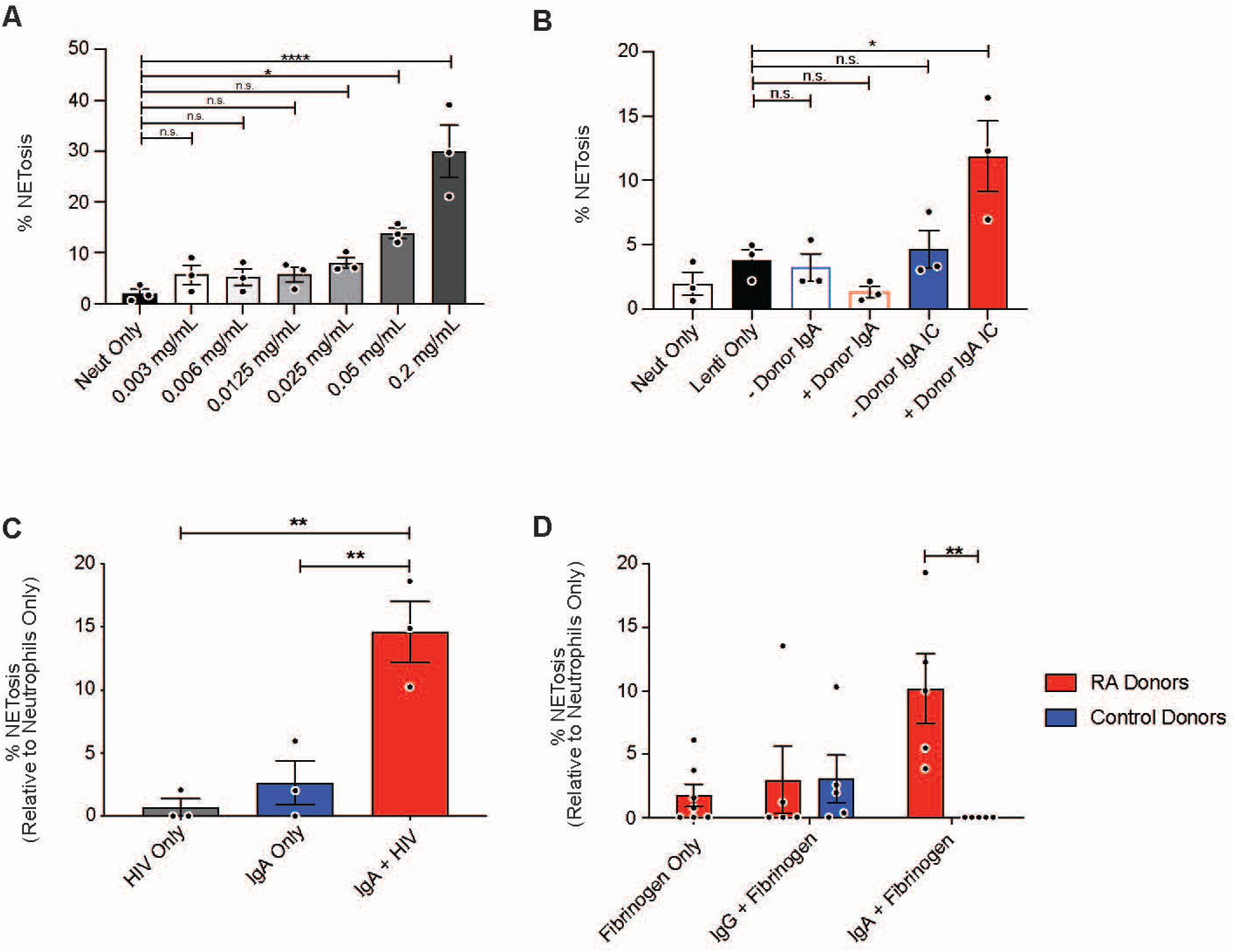
Potentiation of NETosis by IgA ICs is not an IAV-specific phenomenon. (A) Purified SARS-CoV-2 spike pseudotyped lentivirus was titrated onto primary human neutrophils from healthy donors (n = 3) and incubated for 3 hours prior to staining for DNA (DAPI) and neutrophil elastase. (B) Polyclonal IgA was isolated from serum of a convalescent COVID-19 donor and from pre-pandemic donor serum (SARS-CoV-2 seronegative) and incubated with spike pseudotyped-lentivirus to form ICs prior to stimulation of neutrophils isolated from healthy donors (n=3) for 3 hours prior to staining for DNA (DAPI) and neutrophil elastase. (C) ICs were formed with IgA purified from serum of HIV-positive individuals (n=3) and HIV-1 X4 gp120 (HxB2) pseudotyped lentivirus. Neutrophils were stimulated for 3 hours prior to staining for DNA (DAPI) and neutrophil elastase. (D) Cells were stimulated with ICs containing IgA purified from the serum of healthy donors (n=5) or RA patients (n=5) in complex with citrullinated fibrinogen. NETosis was quantified in a blinded manner from 5 fields per condition. Mean and SEM of independent experiments are shown. P-values were determined by one-way ANOVA with Tukey post-hoc test. *, P < 0.05, **, P < 0.01.

We also incubated neutrophils with antibody:HIV ICs which contained HIV-specific IgA isolated from the serum of HIV+ individuals. Following stimulation, a significant increase in NETosis was observed in cells treated with anti-HIV IgA containing ICs (Fig. 2C). Background levels of NETosis wcre observed when cells were treated with either IgA or virus alone. These findings demonstrate that IgA induced NETosis likely happens in the context of many viral infections.

NETs have also been implicated in the pathogenesis of a variety of autoimmune conditions, including rheumatoid arthritis (RA) where they serve as a source of autoantigen ^36,37^. Patients with autoimmune diseases commonly have autoantibodies against NET elements such as histones, DNA, and neutrophil elastase. Here, neutrophils were stimulated with Ab:autoantigen ICs composed of IgA or IgG purified from the serum of RA patients or healthy donors, and recombinant citrullinated human fibrinogen, a common autoantigen in RA ^38^. Induction of NETosis was observed in neutrophils stimulated with IgA:citrullinated fibrinogen immune complexes from RA patients, but not in those stimulated with IgG-containing ICs or ICs generated with antibodies from healthy donors (Fig. 2D). Together, these data demonstrate that potentiation of NETosis is a common property of virus:IgA immune complexes, as well as ICs composed of IgA:autoantigens.

### Induction of NETosis by IgA immune complexes is dependent on FcαRI and independent of TLR signaling

We next assessed whether salivary IgA (sIgA) was capable of inducing NETosis. Whereas monomeric IgA (mIgA) is found predominantly in circulation, secretory IgA is enriched at mucosal surfaces and is generally regarded as an anti-inflammatory antibody. sIgA from saliva and serum-derived mIgA was purified from matched vaccinated donors used to generate ICs with IAV. ICs containing sIgA did not potentiate NETosis, whereas serum-derived mIgA from the same donors was capable of eliciting NETosis, as we had observed previously (Fig. 3A). These results are consistent with previous studies that have demonstrated that the secretory component sterically blocks binding of secretory IgA to FcαRI (CD89) ^39^.

**Figure 3.**
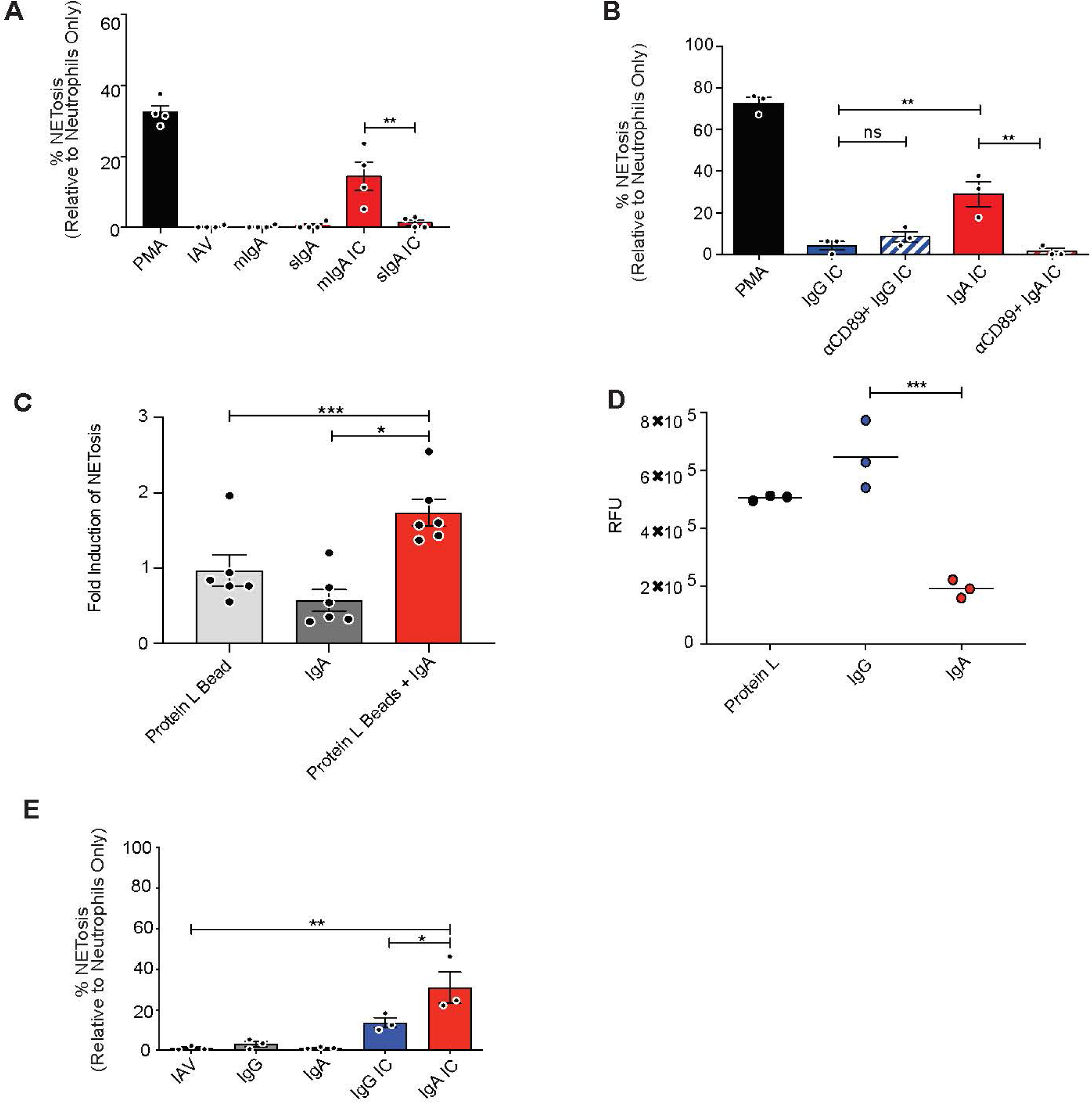
IgA ICs induce NETosis via FcαRI engagement, independently of TLR signaling and phagocytosis. (A) Primary human neutrophils were stimulated for 3 hours with antibody:IAV ICs generated from matched salivary IgA (sIgA) and serum IgA of healthy IAV-exposed donors (n=4). (B) Primary human neutrophils were incubated with an anti-CD89 (FcαRI) antibody prior to stimulation with IgG:IAV or IgA:IAV ICs (n=3) (C) Primary human neutrophils were stimulated with polyclonal IgA, polystyrene beads coated with Protein L, or polystyrene beads coated with protein L and IgA. For all experiments, NETosis was assessed by immunofluorescence microscopy analysis of cells co-stained for DNA (DAPI) and neutrophil elastase. NETosis in stimulated conditions was normalized to untreated cells (n=6). (D) Fluorescent polystyrene beads were coated with protein L, followed by either polyclonal IgG or IgA. Human neutrophils were isolated and incubated with the beads at a 500 beads/cell ratio. After washing, phagocytosis of beads was measured using a SpectraMax i3 plate reader (Molecular Devices) (n=3). (E) Purified Cal/09 was immobilized on glass coverslips prior to the addition of IgG or IgA. Primary human neutrophils were added to wells for 3 hours before being fixed and stained for quantification (n=3). Mean and SEM of independent experiments are shown. Statistical significance was evaluated by one-way ANOVA and Tukey post-hoc test. *, P < 0.05; **, P < 0.01.

Given the observation that sIgA:IAV ICs failed to induce NETosis, we investigated whether mIgA-mediated NETosis was dependent on engagement of FcαRI (CD89). To this end, neutrophils were incubated with a blocking monoclonal anti-CD89 antibody prior to stimulation with IgG:virus or IgA:virus ICs. Blocking with anti-CD89 abrogated induction of NETosis following stimulation with IgA ICs (Fig. 3B), confirming that engagement of FcαRI is required for IgA:virus IC-mediated induction of NETosis.

TLR8 activation has been shown to shift neutrophils from phagocytosis to NETosis in the context of IgG IC-mediated NETosis via FcγRIIA signaling ^32^. We thus set out to determine whether TLR signaling was required for IgA-mediated NETosis induction. TLR8 senses single-stranded RNA and is an important pattern-recognition receptor during RNA virus infection ^40^. Since IAV particles contain RNA, we elected to use a system free from TLR7/8 ligands. To this end, polystyrene beads (roughly equal in number to IAV particles used in previous experiments) were coated with protein L and polyclonal IgA. Protein L binds to the κ light chain of antibodies, leaving the antibody Fc region capable of interacting with FcRs on the cell surface. Following stimulation, IgA:bead ICs induced significant NETosis relative to beads alone (Fig. 3C). This suggests that unlike IgG IC-mediated NETosis, IgA IC-mediated NETosis is likely independent of TLR signaling.

Neutrophils are professional phagocytes, and ADCP is one of the many Fc-mediated effector functions that contribute to their defense against pathogens ^24^. To directly measure whether IgA ICs induced phagocytosis, fluorescent, protein L-coated polystyrene beads were complexed with IgA or IgG prior to incubation with neutrophils. After incubation with beads, cells were washed extensively to remove any beads that had not been phagocytosed. Significantly greater bead uptake was recorded for neutrophils that were exposed to the IgG-opsonized beads compared to those coated with IgA, which actually inhibited phagocytosis relative to protein L-coated control beads (Fig. 3D). This further demonstrates that endosomal TLR activation by viral pathogen-associated molecular patterns (PAMPs) are not required for the potentiation of NETosis by IgA:virus ICs. As further confirmation, instead of using soluble ICs as had been done in previous experiments, IC’s were immobilized on glass coverslips. Consistent with all experiments that had been performed using soluble ICs, significantly higher levels of NETosis were observed when neutrophils were incubated with immobilized IgA:virus ICs relative to immobilized IgG:virus containing ICs (Fig. 3E). Combined, these data suggest that phagocytosis is not required for IgA IC-mediated stimulation of NETosis.

### IgA ICs stimulate NADPH oxidase complex (NOX)-dependent suicidal NETosis

The most common and well-characterized type of NETosis is called “suicidal NETosis”, which results in the death of the cell. More recently, other types of NETosis have been described, including “vital” NETosis ^41^. Suicidal NETosis requires ROS production and occurs between 1-3 hours after stimulation, while vital NETosis does not require the generation of ROS and occurs between 5 and 60 minutes after stimulation ^41^. To determine whether IgA:virus IC-induced NETosis was vital or suicidal, we first performed a time-course experiment following stimulation with PMA, a well characterized stimulant of suicidal/ROS-dependent NETosis, or IgA:IAV ICs for 30, 90 or 180 minutes (Fig. 4A). A significant increase in NETosis was observed following incubation with IgA:IAV ICs for 180 minutes, consistent with suicidal NETosis. Unsurprisingly, PMA – a far more potent stimulant, significantly induced NETosis beginning at 90 min after stimulation (Fig. 4A). Conversely, to inhibit the production of ROS, a small molecule inhibitor of the NOX complex, diphenyleneiodonium chloride (DPI), was pre-incubated with neutrophils prior to stimulation with IgA:IAV ICs. DPI completely inhibited NETosis induced by IgA ICs (Fig. 4B). Together, these observations demonstrate that IgA:virus ICs stimulate suicidal NET release in a NOX-dependent manner.

**Figure 4.**
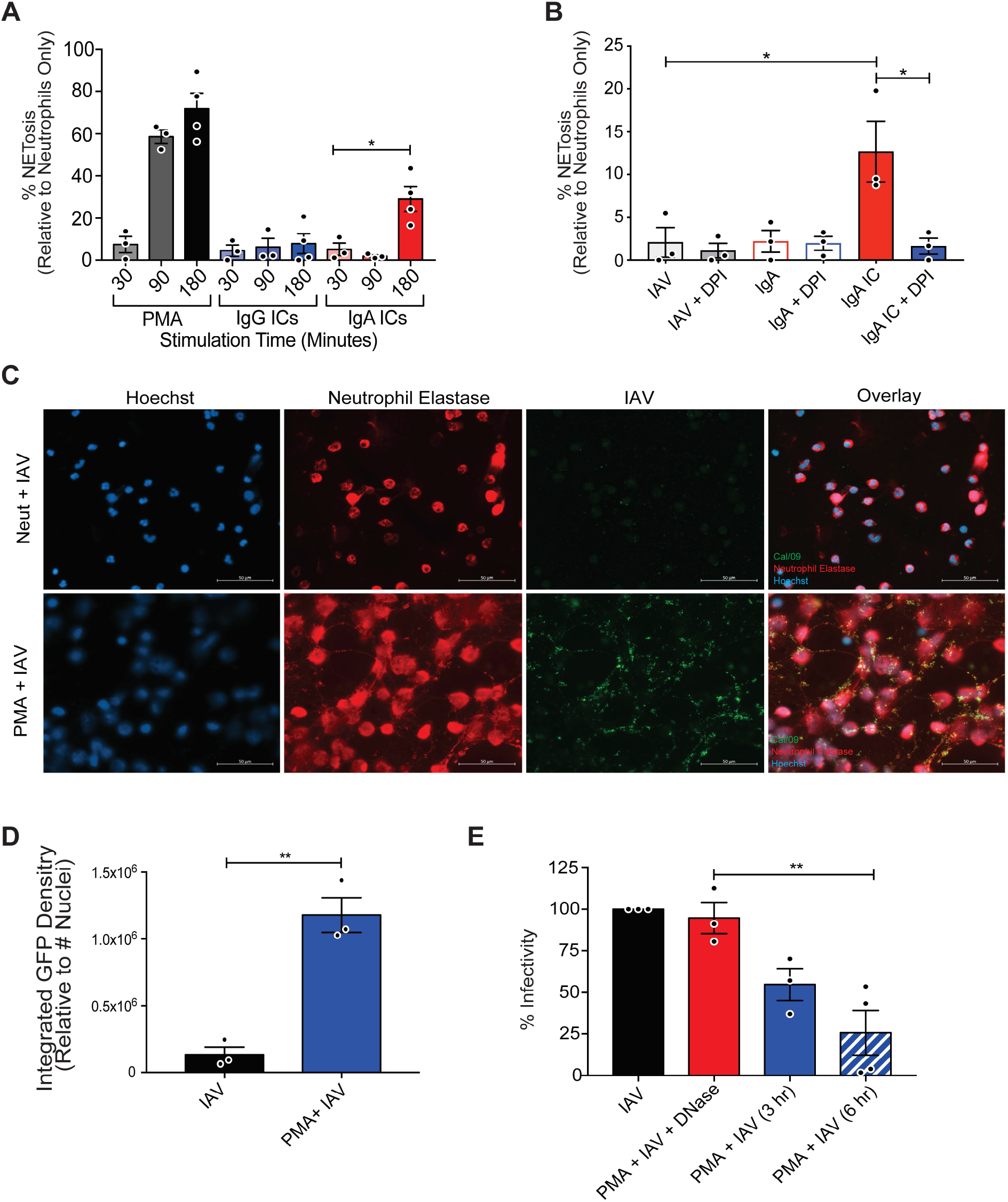
Influenza virus particles are trapped and inactivated by NETs released via suicidal NETosis. (A) Primary human neutrophils were stimulated with PMA, IgG:IAV, or IgA:IAV ICs for 30, 90 and 180 minutes prior to fixation and staining for DNA (DAPI) and neutrophil elastase to quantify NETosis. (B) Primary human neutrophils were incubated with DPI, a NOX inhibitor, prior to 3 hours of simulation with IgA:IAV ICs (n=3). (C) Neutrophils were stimulated with PMA for 90 minutes before the addition 105 PFU/ well of IAV. Virus was incubated with the NETs for 3 hours and then fixed and stained with anti-hemagglutinin antibodies (6F12), DNA (DAPI) and neutrophil elastase. Immunofluorescence microscopy was used to measure the co-localization of viral particles (green) with NETs composed of DNA (blue) coated with neutrophil elastase (red) (n=3) (D) Quantification of the raw integrated GFP density was measured using ImageJ and normalized to the number of cells per field. (E) Neutrophils were stimulated with PMA prior to the addition of an IAV expressing an mNeon reporter. Virus was incubated on intact NETs or NETs that had been digested with DNase (n=3). Contents of wells were collected and MDCK cells were infected for 8 hours to measure residual infectivity. Mean and ± SEM are shown. Statistical significance was evaluated using one-way ANOVA with Tukey post-hoc test. *P < 0.05; **P < 0.01.

### Virus particles are trapped and inactivated by NETs

In the context of bacterial infections, NETs exert antimicrobial activity by trapping and killing bacteria with antimicrobial effector proteins associated with NETs. We thus set out to determine whether NETs were similarly capable of trapping and inactivating virus. Neutrophils were either left unstimulated or were treated with PMA to induce suicidal NETosis (virus containing ICs were not used to avoid the confounding issue of having viruses present during induction of NETosis). IAV was then incubated in wells of stimulated or unstimulated neutrophils, and unbound virus was washed away. Using immunofluorescence microscopy, we observed that IAV particles become trapped in NETs induced following stimulation with PMA (Fig. 4C). Using ImageJ software, we quantified GFP pixel density and normalized this to the number of cells (and/or NETs) per field. Consistent with the stark visual contrast observed in the images, significantly more virus was associated with PMA-stimulated neutrophils that had undergone NETosis than unstimulated neutrophils (Fig. 4D).

To test whether IAV was inactivated after being trapped in NETs, we used an mNeon reporter virus ^42^. IAV-mNeon was incubated with unstimulated neutrophils, PMA-stimulated neutrophils that had undergone NETosis, or PMA-stimulated neutrophils treated with DNase to digest NETs. DNase digestion specifically allowed us to test whether being trapped in a NET was necessary for inactivation, or whether factors released by neutrophils during NETosis were alone sufficient to inactivate IAV (Supplementary Fig. 1, Fig. 4E). After 3h or 6h incubation, viral media was collected from all wells incubated on Madin Darby Canine Kidney cells (MDCKs) to quantify remaining infectious virus. Incubation of virus with PMA-stimulated neutrophils that had undergone NETosis significantly reduced infectivity after 3 h and 6 h incubation. Interestingly, digestion of NETs produced by PMA-simulated cells with DNase prior to addition of virus had no significant impact of infectivity – suggesting that physical contact with NETs is required for inactivation and that soluble factors released during the process of NETosis alone are not sufficient to mediated inactivation (Fig. 4E). Taken together, these data demonstrate the viruses can be trapped and inactivated by NETs.

## DISCUSSION

NETosis has been most extensively studied as an anti-pathogen immune response in the context of bacterial infections ^43^. However, accumulating evidence suggests that NETs have antiviral activity, but can also contribute to the pathogenesis of viral disease in certain circumstances ^8–11,44,45^. While pathogens like viruses and bacteria can trigger NETosis directly as an innate immune mechanism, there is also an important intersection of neutrophils/NETs and the adaptive immune response, since neutrophils express Fc receptors capable of recognizing both soluble ICs and antibody-bound cells. Here, we show that IgA significantly lowers the amount of virus required to trigger NETosis.

In the context of IgG, immobilized ICs have been reported to stimulate NETosis via FcγRIIA. Soluble ICs were primarily phagocytosed, but could be shifted to stimulate NETosis upon TLR7/8 activation, which resulted in furin-mediated cleavage and shedding of the FcγRIIA N-terminus – inhibiting further phagocytosis ^32^. We observed that IgA ICs did not simulate phagocytosis, but rather preferentially induced NETosis, even in the absence of TLR activation. While IgG ICs could stimulate NETosis, the induction of NETosis was notably more pronounced upon stimulation of neutrophils with IgA ICs.

In the context of IAV, bnAbs that bind to the conserved HA stalk domain have become a major focus for the development of “universal” influenza virus vaccines and monoclonal antibody prophylactics/therapeutics. Although bnAbs are relatively weak neutralizers of IAV, they confer protection *in vivo* by potent induction of Fc-dependent effector functions ^17–19,22,24,26,27^. The ability of bnAbs to elicit potent effector functions (relative to conventional neutralizing antibodies that bind to the HA head domain) relies on a unique reciprocal contact model whereby Fc receptors of immune effector cells bind to the Fc domain of bnAbs bound to HA on target cells, while HA expressed on target cells in turn binds to sialic acid residues of the effector cell ^26,27^. However, almost everything that is known about the function of bnAbs has been studied in the context of IgG. Of the other immunoglobulin isotypes, IgA plays a particularly important role in protection against mucosal viruses. Indeed, local IgA responses correlate with protection offered by live-attenuated influenza virus vaccines ^46–48^. A recent Phase I trial of a chimeric HA universal vaccine candidate reported potent induction of IgA bnAbs after vaccination – further highlighting the urgent need to understand how antibodies of this isotype contribute to protection ^20,49^. Here, we show that consistent with prior studies, bnAbs are primarily responsible for induction of FcαRI-dependent NETosis, likely because these antibodies also promote the reciprocal binding events between IgA:FcαRI and HA:sialic acid described above.

Importantly, the ability of IgA ICs to potentiate NETosis was widespread across several different viruses including IAV, lentiviruses pseudotyped with SARS-CoV-2 S protein, and HIV. Indeed, this phenomenon could also be recapitulated with IgA-coated beads and extended beyond the context of infectious diseases to ICs composed of IgA from RA patients in complex with citrullinated fibrinogen – a common RA autoantigen. Our findings support previous work from the van Egmond laboratory demonstrating that IgA ICs isolated from synovial fluid of RA patients also induce NETosis ^36^. Previous work by our group has shown that upon exposure to IgG:IAV ICs, neutrophils undergo ADCP and potently induce ROS. Inhibition of phagocytosis with cytochalasin D almost completely abolished ROS induction by IgG:IAV ICs. In contrast, IgA:IAV ICs were able to stimulate ROS even when phagocytosis was inhibited ^24^. Those observations are in line with the data presented herein showing that IgA:ICs induced neutrophils to undergo ROS-dependent suicidal NETosis in a phagocytosis-independent manner.

In serum, IgA is present at concentration of ∼ 82-624 mg/dL, whereas IgG is found at ∼ 694-1803 mg/dl ^50^. In the context of HIV, mixed IgG/IgA:HIV ICs generated using the gp41-specific bnAb 2F5 cooperatively triggered ADCC of HIV-infected cells by monocytes, but did not act cooperatively to induce ADCP ^33,51^. Likewise, we observed no cooperativity in the induction of NETosis when IgA and IgG were combined at a 1:1 ratio, or at physiological ratios. These results suggest that signaling downstream of FcγRs and FcαRI lead to distinct effector outcomes in monocytes and neutrophils.

While ICs composed of serum-derived IgA and monomeric monoclonal IgA could both potentiate NETosis, secretory IgA purified from saliva could not. This is consistent with prior studies that have demonstrated that the secretory component sterically interferes with binding to FcαRI and suggests that IgA-stimulated NETosis is unlikely to occur in the airways where secretory IgA is enriched, but instead would be expected to take place primarily in tissues and vasculature ^52^. The data presented here demonstrate that NETs can both trap and inactivate virus. This suggests that they may have a protective antiviral function. We speculate that in individuals who lack virus-specific IgA, the high concentrations of virus needed to stimulate NETosis might exacerbate inflammation and potentiate disease, as has been observed for those with COVID-19 ^14–16,35^. However, individuals with pre-existing immunity – such as that conferred by vaccines – low levels of IgA-induced NETosis might help to trap and inactivate virus early in infection, thereby limited virus spread and progression to severe disease.

In summary, we report a new antiviral effector function mediated by virus:IgA ICs. The mechanism through which virus:IgA ICs stimulate NETosis is distinct from, and considerably more potent than virus alone. Since mice do not express an FcαR, it will be important to develop alternative models for *in vivo* studies to determine when IgA:virus IC-mediated NETosis may be protective, and when it may exacerbate disease.

## MATERIALS AND METHODS

### Human Serum and Blood Samples

Human blood samples used to isolate serum antibodies were obtained with permission from consenting male and female IAV-vaccinated donors, SARS-CoV-2 infected donors, HIV-positive individuals, and RA patients. Human blood for neutrophil isolations were collected with permission from consenting healthy male and female donors. All protocols involving human samples were approved by the Hamilton Integrated Research Ethics Board and the Western Research Ethics Board. Blood was collected into Ethylenediamine tetra-acetic acid (EDTA) coated tubes (BD Vacutainer).

### Neutrophil Isolation

Neutrophils were isolated from the peripheral blood of healthy male and female donors by density gradient centrifugation as described previously ^24^. Briefly, 3 mL of room temperature Histopaque 1119 (Sigma-Aldrich) was added to a 15 mL falcon tube, followed by gentle addition of 3 mL of Histopaque 1077 (Sigma-Aldrich). 6 mL of blood was layered on top and samples were centrifuged at 930 x g for 30 minutes at room temperature (RT) with no deceleration in an Allegra X-12R centrifuge (Beckman Coulter). The neutrophil layer was collected between the Histopaque layers and diluted in 4°C PMN buffer (0.5% BSA, 0.3 mM EDTA in Hank’s balanced salt solution (Sigma-Aldrich)) to a total volume of 50 mL. PMNs were then centrifuged at 450 x g for 5 minutes at RT. The Supernatant was discarded, and the cell pellet re-suspended by flicking the tube. To lyse red blood cells, 3 mL of ACK (ammonium-chloride-potassium) lysis buffer (8.3 g/L NH_4_Cl, 1 g/L KHCO_3_, 0.05 mM EDTA, in sterile distilled H_2_O) was added to the PMNs and incubated for 3 minutes with agitation every 30 seconds. The PMNs were diluted in 30 mL of PMN buffer and centrifuged at 450 x g for 5 minutes at RT, followed by one additional wash.

### Antibody Purification

Heat-inactivated human serum was diluted 1:10 in phosphate buffered saline (PBS) and applied to a gravity polypropylene flow column (Qiagen) containing 1 mL of Protein G-sepharose resin (Invitrogen) to purify IgG. Flow through sera was then applied to a gravity flow column containing 1 mL Peptide M-sepharose resin (InvivoGen) to purify IgA. Columns were washed with two column volumes of PBS. IgG and IgA were eluted with 0.1 M glycine-HCl buffer (pH 2.7) into 2 M Tris-HCl neutralizing buffer (pH 10). Antibodies were concentrated and re-suspended in PBS using 30 kDa cutoff Macrosep Advanced Centrifugal Devices (Pall Corporation). To purify monoclonal antibodies, clarified cell culture supernatants were applied directly to Protein G-sepharose columns prior to washing and elution.

### Monoclonal Antibodies

The variable light and heavy chain sequences of KB2 and 29E3 antibodies ^30,53^ were cloned into cloned into pFuse vectors (pFUSE-hIgG1-Fc2 and pFUSE2ss-CLIg-hK, Invivogen). KB2 binds to the stalk domain of H1 viruses, while 29E3 antibody is specific to the head domain of A/California/04/09 (Cal/09). HEK293T cells co-transfected with pFUSE plasmids according to manufacturer’s recommendations and were subsequently purified from supernantants using Protein G-sepharose columns, as described above.

### Cells and Viruses

Madin Darby Canine Kidney (MDCK) cell were grown in Dulbecco modified Eagle medium (DMEM) containing 10% fetal bovine serum (FBS) (Gibco), 2 mM L- and 100 U/mL penicillin-streptomycin (Thermo Fisher). At 100% confluency MDCK cells were infected for one hour with A/California/04/2009 H1N1 (kind gift of Dr. Peter Palese, Icahn School of Medicine at Mount Sinai, New York, NY) in 1x minimum essential medium (MEM, Sigma Aldrich) supplemented with 2 mM L-glutamine, 0.24% sodium bicarbonate, 20 mM HEPES (4-(2-hydroxyethyl)-1-piperazineethanesulfonic acid), MEM amino acids solution (Sigma Aldrich), MEM vitamins solution (Sigma Aldrich), 100 U/mL penicillin-streptomycin (Thermo Fisher), and 0.42% bovine serum albumin (Sigma Aldrich). Cells were then washed with PBS and media was replaced. Cells were left for 72 hours and supernatant was collected. A/ Puerto Rico/8/1934/ H1N1-mNeon (PR8-mNeon, which was a kind gift from the laboratory of Dr. Nicholas Heaton (Duke University, Durham, NC) ^42^, was propagated in 10-day-old embryonated chicken eggs as per standard protocols (WHO, 2011).

### Influenza Virus Purification

Clarified supernatants from IAV-infected MDCK cells were layered on top of 8 mL 20% sucrose (Bioshop) in NTE buffer (0.5 M NaCl, 10 mM Tris-HCl, 1 mM EDTA, pH 7.5) inside Ultra-Clear Ultracentrifuge tubes (Beckman Coulter). Samples were spun at 76,650 x g for 2 hours at 4° C inside a SW 32i rotor using an Optima L-90K Ultracentrifuge (Beckman Coulter). Purified virus was quantified using a bicinchoninic acid assay (BCA) Protein Assay Kit (Pierce Biotechnology) according to the manufacturer’s instructions, and by hemagglutination assay.

### Psuedotyped Lentivirus Production

HIV-1 X4 gp120 pseudotyped lentiviruses were prepared described previously ^55^. Briefly, HEK293T cells were cultured in DMEM supplemented with 10% fetal bovine serum, L-glutamine, 100U/ml penicillin and streptomycin and maintained in 5% CO_2_ at 37^°^C. Briefly, 5 ×10^5^ cells were seeded onto 6-well plates in one day prior to transfection. Cells were co-transfected on the next day at 70-80% confluency with pLenti-CMV-GFP-Puro (1.5µg) along with pEnv_HxB_ (0.5µg) and psPAX2 (1µg) plasmids. Medium was changed 24h post-transfection. Supernatant was then harvested, filtered with a 0.22 micron filters (Millipore) and titered as described previously ^55^. The virus was stored at −80°C until use.

SARS-CoV-2 S protein pseudotyped lentiviruses were produced as described by Crawford et al.^56^ and the following reagents were obtained through BEI resources, NIAID, NIH: SARS-Related Coronavirus 2, Wuhan-Hu-1 Spike-Pseduotyped Lentiviral Kit, NR-52948. In brief, HEK293T cells seeded in 15cm dishes at 1.1×10^7^ cells/mL in 15 mL of standard DMEM. 16-24 hours post seeding, cells were co-transfected with HDM-nCoV-Spike-IDTopt-ALAYT, pHAGE-CMV-Luc2-IRES-ZsGreen-W (BEI catalog number NR-52516), HDM-Hgpm2 (BEI catalog number NR-52517) HDM-tat1b (BEI catalog number NR-52518) , pRC-CMV-Rev1b (BEI catalog NR-52519). 18-24 hours post transfection the media was replaced with full DMEM. 60 hours post transfection, the supernatant was collected and filtered with a 0.45 µm filter and stored at −80 degrees. For purification, 40 mL of supernatant was concentration by spinning at 19, 400 rpm for 2 hours. The resulting pellet was resuspended in 400µl of HBSS, followed by 15 mins of continuous vortex at RT. Protein concentration was confirmed by BCA.

### Coating Polystyrene Microspheres with Protein L and Polyclonal Antibody

Fluorescent carboxylate microspheres, 0.5µm (Polysciences) were coated with Protein L (Thermo Scientific), followed by polyclonal IgA or IgG. The polystyrene microspheres were first washed with 1X PBS and centrifuged at 13 523 x g. The PBS wash was repeated and then the microspheres were incubated at RT with 750 µg of Protein L for 4 hours with gentle mixing. Following another PBS wash, 300 µg of polyclonal IgA was added to the microspheres and left to incubate at RT with gentle mixing overnight. Following this incubation, the microspheres were centrifuged at 13 523 x g for ten minutes, and the resulting pellet was resuspended in 1 mL of PBS and incubated with for 30 minutes with gentle mixing at RT. Following the final incubation, the microspheres were centrifuged at 13 523 x g for 5 minutes and resuspended in 500 µL of PBS.

### Neutrophil Stimulation with Soluble ICs

15 mm glass coverslips were placed inside wells of a sterile 24-well plate and 4.0×10^5^ PMNs were added to each well and allowed to settle for 1 hour. For IAV:polyclonal Ab stimulations, mixtures of 25 µg Cal/09 (2^10^ HAU) and 50 µg/mL polyclonal IgG or IgA antibody were incubated for 30 minutes at 4°C before addition to PMNs. ICs containing monoclonal HA stalk (KB2) or head-binding (29E3) antibodies were generated at a 2:1 ratio of antibody to virus (100 µg/ml and 50 µg/ml, respectively) and allowed to incubate for 30 minutes at RT prior to stimulation of PMNs. To test the ability of ICs generated with salivary IgA to stimulate NETosis, matched salivary IgA and serum IgA was purified from the saliva of four healthy donors using peptide M columns. ICs containing 100 µg/well of serum derived monomeric IgA or salivary IgA and 50 µg/ well of Cal/09 were allowed to form by incubation at RT 30 minutes. HIV-specific ICs were generated by purifying IgA from the serum of 3 HIV-1 positive donors. ICs were formed by incubating 100 µg/mL of polyclonal IgA and 50 µg/mL of HIV-1 gp120 pseudotyped lentiviruses for 30 minutes at room temperature. SARS-CoV-2 ICs were generated using antibodies purified from the convalescent sera of an individual who had been infected with SARS-CoV-2. ICs were formed by incubating 100 µg/mL of polyclonal IgA and 12.5 µg/mL of pseudotyped spike lentiviruses for 30 minutes at room temperature. For RA samples, immune complexes were formed by incubating 50 µg/well of citrullinated human fibrinogen (Cayman Chemicals) with 100 µg/well of polyclonal IgA or IgG for 30 minutes at RT. Stimulation of IgA-coated beads was performed by incubating neutrophils with 5.0 × 10^8^ beads. Antibodies were purified from the sera donors diagnosed with RA or healthy donors as described above. Antibodies/viruses/ICs were then incubated with PMNs for 3 hours at 37°C before being fixed with 3.7% paraformaldehyde (PFA) (Pierce Protein Biology) prior to staining and imaging.

### Immobilized IC Assay

Purified virus was plated on 15mm sterile coverslips in a 24-well plate at 2µg/mL and incubated at 37°C for 18 hours. Wells were washed twice with PBS and 250 µg of either polyclonal IgA or IgG was added for 30 minutes at 37°C. Wells were washed twice with PBS prior to the addition of 4.0 ×10^5^ PMNs per well. PMNs were incubated for 3 hours at 37°C, fixed and stained as described above.

### FcαRI Blockade

15 mm glass coverslips were placed inside wells of a sterile 12-well plate and 4.0×10^5^ PMNs were added to each well in a total volume of 500 µL and allowed to settle for 1 hour. To block FcαRI, 20 µg/mL of mouse anti-human CD89 antibody (AbD Serotec) was added to neutrophils for 20 minutes at 4°C. PMNs were then stimulated with various conditions for 3 hours at 37°C before being fixed with 3.7% paraformaldehyde (PFA) and stored at 4°C until staining.

### Fluorescence Microscopy and Quantification of NETosis

Cells were fixed with 3.7% PFA (Pierce Protein Biology) at 4°C, washed in PBS three times and then permeabilized using 0.5% Triton X-100 (Thermo Scientific) in PBS-T. Fixed and permeabilized cells were then blocked for 30 minutes at RT in blocking buffer (10% FBS in PBS-T). Cells were incubated with primary rabbit anti-neutrophil elastase antibody (Abcam) at a 1:100 dilution for one hour at room temperature. Coverslips were washed with PBS three times and then incubated with Alexa Fluor 488-conjugated donkey anti-rabbit antibody (Molecular Probes) diluted as per manufacturer’s recommendation (2 drops/mL) for 1 hour at RT, protected from light. Coverslips were then washed with PBS three times. 1 µg/mL Hoechst 33342, trihydrochloride, trihydrate (Life Technologies) was incubated for 5 minutes, at RT, protected from light. Cells were washed with PBS three times and coverslips were mounted onto glass slides in EverBrite Mounting Medium (Biotium). Cells were imaged using an EVOS FL microscope (Life Technologies). 5 random fields per condition were captured at 20x magnification. NETosis was quantified by counting cells which had decondensed chromatin colocalized with neutrophil elastase. % NETosis was expressed as number of cells that had undergone NETosis / number of total cells.

### Influenza Viral Particle Trapping and Inactivation in NETs

Sterilized glass coverslips were placed in a 24 well plate, and neutrophils at 4.0×10^5^ cells/well were allowed to settle for 1 hour prior to stimulation. Neutrophils were stimulated with PMA for 3 hours at 37 °C, 5 % CO_2_. Cal/09 at 10^5^ PFU/mL was then allowed to settle on the pre-formed NETs for 3 hours at 37°C, following this incubation cells were fixed with 3.7% PFA (Pierce Protein Biology). To stain coverslips for immunofluorescent imaging coverslips were treated in the same was as previously described. Primary antibodies used included primary rabbit anti-neutrophil elastase antibody (Abcam, 1:100 dilution), 6F12 generated from in house-hybridomas at 1 ug/mL. Secondary antibodies included Alexa Fluor 488 conjugated donkey anti-mouse antibody (Molecular Probes, 1:4000) and Alexa Flour 594 donkey anti-rabbit (Molecular Probes, 1:4000). Coverslips were incubated with 1 µg/mL Hoechst 33342, trihydrochloride, trihydrate (Life Technologies) to probe for DNA. Cells were visualized and imaged using GFP (Ex 470 nm/Em 525 nm) and DAPI (Ex 360 nm/Em 447 nm), Texas Red (Ex 585/ Em 624) color cubes in the EVOS FL microscope (Life Technologies). To evaluate inactivation, 10^5^ PFU/mL of PR8 mNeon was incubated with PMA stimulated neutrophils for 3-6 hours. 25 units/mL of DNaseI (Thermo Fisher) was added to PMA stimulated neutrophils and was allowed to incubate for 90 minutes to digest NETs. Samples were collected and stored at −80 until further use. Prior to virus quantification, MDCK cells were seeded in 24-well plates and used when 90% confluent. Sample was diluted 1:10 in 1x minimum essential medium (MEM, Sigma Aldrich) supplemented with 2 mM L-glutamine, 0.24% sodium bicarbonate, 20 mM HEPES (4-(2-hydroxyethyl)-1-piperazineethanesulfonic acid), MEM amino acids solution (Sigma Aldrich), MEM vitamins solution (Sigma Aldrich), 100 U/mL penicillin-streptomycin (Thermo Fisher), and 0.42% bovine serum albumin (Sigma Aldrich), before being added to cells. After 1 hour this was replaced with DMEM containing 10% FBS (Gibco), 2 mM L-glutamine and 100 U/mL penicillin-streptomycin (Thermo Fisher). The number of fluorescent cells was assessed 12 hours post-infection. Cells were fixed with PFA and incubated with 1 µg/mL Hoechst 33342, trihydrochloride, trihydrate (Life Technologies). 5-fields per condition were taken on the EVOS FL microscope, and % infectivity was determined as the number of infected cells / the total number of cells.

### Phagocytosis Assay of Polyclonal Antibody-Coated Microspheres

This protocol was performed as previously described ^24^. Briefly, fluorescent carboxylate microspheres 0.5µm (Polysciences) were coated with protein L and polyclonal IgA or IgG and were incubated with neutrophils at a 500:1 ratio at 37°C for 15 minutes with gentle mixing. This was followed by centrifugation at 930 × g for 10 minutes. Cells were washed twice with PBS before being plated in a 96-well plate. Fluorescence was measured with the SpectraMax i3 plate reader at 526nm (Molecular Devices).

### NOX Assay

Neutrophils were purified as described above and allowed to settle on glass coverslips for 1 hour at 37 °C. While settling, neutrophils were incubated with 20 µm of Diphenyleneiodonium chloride (DPI) (Sigma-Aldrich) a neutrophil NADPH oxidase inhibitor. Neutrophils were then stimulated with IgA:IAV ICs or PMA (0.1 mg/mL, Sigma-Aldrich) as a positive control and DPI was maintained in the media. Cells were then fixed with 3.7% paraformaldehyde (PFA) (Pierce Protein Biology) and stored at 4°C until staining and imaging.

### Statistics

Graphs and statistical analyses were generated using Graphpad Prism v9 (Graphpad Software, San Diego, CA). A *P* value of < 0.05 was considered to be significant across all experiments.

## FIGURE LEGENDS

**Supplementary Figure 1. NETs digested with DNase**. Neutrophils were isolated and stimulated with IgA:IAV IC’s or PMA for 3 hours. DNase was added at 25 units/mL and allowed to incubate for 90 min prior to fixation and staining.

## ACKNOWLEDGEMENTS

This work was funded by grants from the Canadian Institutes of Health Research (CIHR) (M.S.M.), the Weston Family Microbiome Initiative (M.S.M.), The Lung Association/Ontario Thoracic Society Grants-in-Aid (M.S.M.), and the Michael G. DeGroote Institute for Infectious Disease Research (M.S.M.). M.S.M. was also supporting, in part, by a CIHR New Investigator Award and an Ontario Early Researcher Award (ERA). H.D.S. was supported, in part, by a CIHR Master’s Award and an Ontario Graduate Scholarship. The authors thank Dr. Joe Mymryk for critical reading of the manuscript and helpful suggestions.

## AUTHOR CONTRIBUTIONS

M.S.M. and C.E.M. conceived project. H.D.S., D.G, A.P., J.C.A., K.E.N., and M.A.Z. performed experiments and analyzed data. C.K., E.C. and D.M.E.B provided critical reagents and experimental input. H.D.S. and M.S.M. wrote the manuscript with input from all authors.

