## Supplementary Figure 1 for "IgA Potentiates NETosis in Response to Viral Infection"

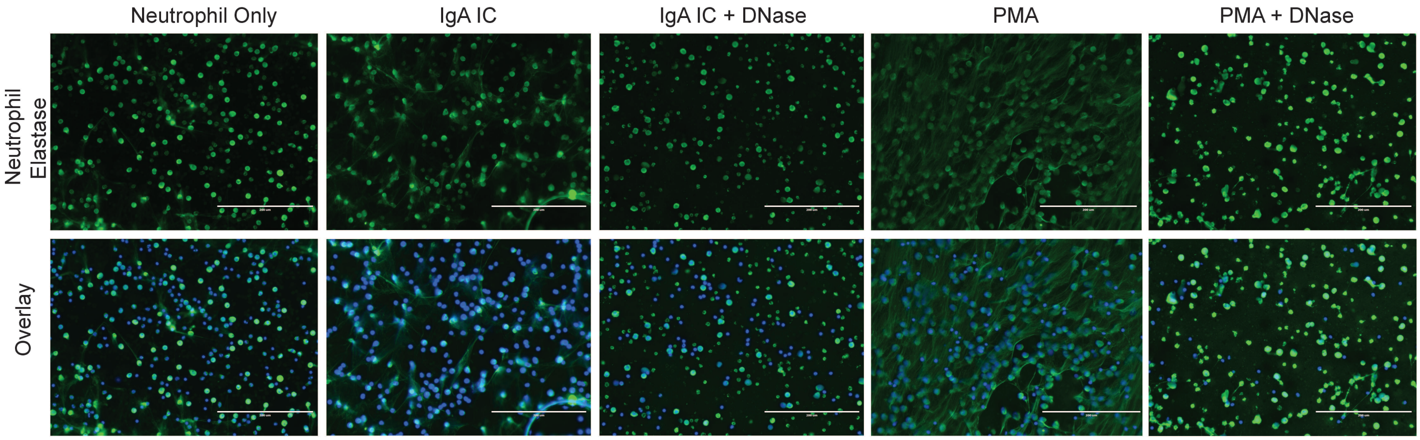


**Supplementary Figure 1. NETs digested with DNase.** Neutrophils were isolated and stimulated with IgA:IAV IC's or PMA for 3 hours. DNase was added at 25 units/mL and allowed to incubate for 90 min prior to fixation and staining.
